# Design, Mutate, Screen: High-throughput creation of genetic clocks with different period-amplitude characteristics

**DOI:** 10.1101/2021.06.22.449269

**Authors:** Andrew Lezia, Nicholas Csicsery, Jeff Hasty

**Author notes:** Correspondence and requests for materials should be addressed to J.H. Authors contributed equally to this work (Andrew Lezia, Nicholas Csicsery).

## Abstract

Directed evolution has become an invaluable tool in protein engineering and has also greatly influenced the construction of synthetic gene circuits. The ability to generate diversity at precise targets for directed evolution approaches has improved vastly, allowing researchers to create large, specific mutant libraries with relative ease. Screening approaches for large mutant libraries have similarly come a long way, especially when the desired behavior can easily be tested for with static, single time-point assays. For more complex gene circuits with dynamic phenotypes that change over time, directed evolution approaches to controlling and tuning circuit behavior have been hindered by the lack of sufficiently high-throughput screening methods to isolate variants with desired characteristics. Here we utilize directed mutagenesis and multiplexed microfluidics to develop a workflow for creating, screening and tuning dynamic gene circuits that operate at the population level. Specifically, we create a mutant library of an existing oscillator, the synchronized lysis circuit, and tune its dynamics while uncovering principles regarding its behavior. Lastly, we utilize this directed evolution workflow to construct a new synchronized genetic oscillator that exhibits robust dynamics over long time scales.

---

Directed evolution has solidified its role as an invaluable tool for biologists to engineer proteins with new or improved functions. For example, directed evolution has been used to develop enzymes that carry out non-naturally occurring reactions^1^, improved fluorescent proteins^2, 3^, and orthogonal sets of RNA polymerase-promoter pairs^4^. While the process of purposefully generating genetic diversity and selecting for desired characteristics stemmed from the field of protein engineering, synthetic biologists have readily acknowledged its potential for creating functional gene circuits^5, 6^. In one of the first examples of directed evolution applied to synthetic biology, Yokobayashi et. al. optimized a genetic circuit composed of two sequential logic gates^7^. Since then, design-then-mutate approaches for synthetic gene circuits have blossomed, with many groups employing this strategy for applications such as optimizing biosynthesis pathways to maximize yield of high-value chemicals^8^. In one recent example, researchers developed a workflow for directed evolution of genetic switches in yeast^9^. Using their platform, they were able to find genetic switches with increased sensitivity and signal-to-noise ratio and use them to improve a caretonoid biosynthesis pathway.

The two key features of any successful directed evolution pipeline are the creation of a population of variants and the selection for desired traits among these variants^10^. Methods to create large, targeted mutant libraries for directed evolution have improved vastly over the past decade allowing researchers to simultaneously mutate multiple genetic targets at once^11, 12^, use host organisms to mutate the desired target *in vivo*^13–15^, and rapidly assemble many pieces of DNA in single reactions^16, 17^. Site directed mutagenesis (SDM) techniques in particular have made the creation of precisely-targeted mutant libraries, easy, inexpensive, and fast^18^.

Similarly, there have been considerable advances in screening and selection strategies for directed evolution, especially when applied to protein engineering^19–21^. Strategies for screening and selecting for desired variants in synthetic gene circuit libraries have also grown considerably, but many of these strategies have focused on gene circuits that do not exhibit complex, dynamic behaviors that change over time^22^. Massively multiplexed microfluidic platforms, like those developed by our group, have been used to screen strain libraries for some dynamic characteristics such as the first-order response to environmental perturbations, but have yet to be employed for engineered gene circuits with free-running dynamic behavior^23^. For these dynamic gene circuits, such as synchronized gene oscillators, the use of directed evolution to tune and improve their function has been limited by the lack of quantitative, high-throughput methods to screen dynamic circuit parameters such as period and amplitude. One group recently developed a method to isolate single-cells after time-lapse microscopy and used this technology to screen large libraries of single-cell oscillator circuits^24^. This technology addressed a gap in the field of high-throughput screening of single-cell dynamic gene circuits, but there is still a need for platforms that allow screening of population-level gene circuits such as synchronized oscillators that rely on quorum sensing for cell-cell communication.

In this work, we utilize a multi-strain microfluidic device to simultaneously culture multiple engineered strains of *E. coli* harboring variants of synthetic oscillator circuits. Specifically, we develop a workflow for quantitatively screening libraries of dynamic gene circuits. We use this workflow to tune the dynamics of an existing oscillator, the synchronized lysis circuit, and uncover new principles regarding its regulation. Additionally, we develop a new synchronized gene oscillator and demonstrate how we are able to improve the dynamics of this new circuit by combining computational modeling with our screening pipeline. The final oscillator we developed exhibits robust and tunable oscillations over long time scales. Overall, this work demonstrates an approach for the directed evolution of dynamic gene circuits and illustrates how multi-strain microfluidics can vastly improve the ability to screen for dynamic gene circuit phenotypes at the population level.

## 1 Results

### Overview of directed evolution approach for synthetic oscillator creation and tuning

We sought to develop and utilize a system for constructing and tuning gene circuits with dynamic phenotypes by directed mutagenesis and screening (Fig. 1). In this work, we focus on the tuning and creation of oscillator circuits in *E. coli* that are synchronized at the population level as they: 1) exhibit complex, time-varying phenotypes that can be difficult to predict and monitor, 2) have many dynamic parameters that can be tuned via mutagenesis (e.g. period, amplitude, and prominence), and 3) are increasingly being tested for real-world applications. We begin by creating targeted mutant libraries of a genetic circuit using site-directed-mutagenesis (SDM). We utilize deterministic modeling of circuit dynamics to help guide the choice of circuit elements to mutate for library creation. Following circuit library creation and transformation into *E. coli*, we screen circuits for interesting phenotypes in both well plate-based batch culture and microfluidic-based continuous culture. Batch culture approaches for gene circuit library screening allow for simple and rapid screening of many variants for significant phenotype differences, but are insufficient for carefully screening for dynamic phenotypes that are only seen in a continuous culture environment where the metabolic state of the growing cell population is relatively constant. Thus, after an initial library screen in 96 well-plates, we deploy a high-throughput multi-strain microfluidic device to further screen interesting oscillator library members for dynamic phenotype differences. Specifically, we are able to pattern up to 48 unique strains of *E. coli* to distinct positions on the microfluidic device and grow them continuously for mutliple days while maintaining precise control of the media composition and flow rate. The high spatio-temporal resolution data from variants can be used to improve circuit models and inform design considerations for relevant applications.

**Figure 1:**
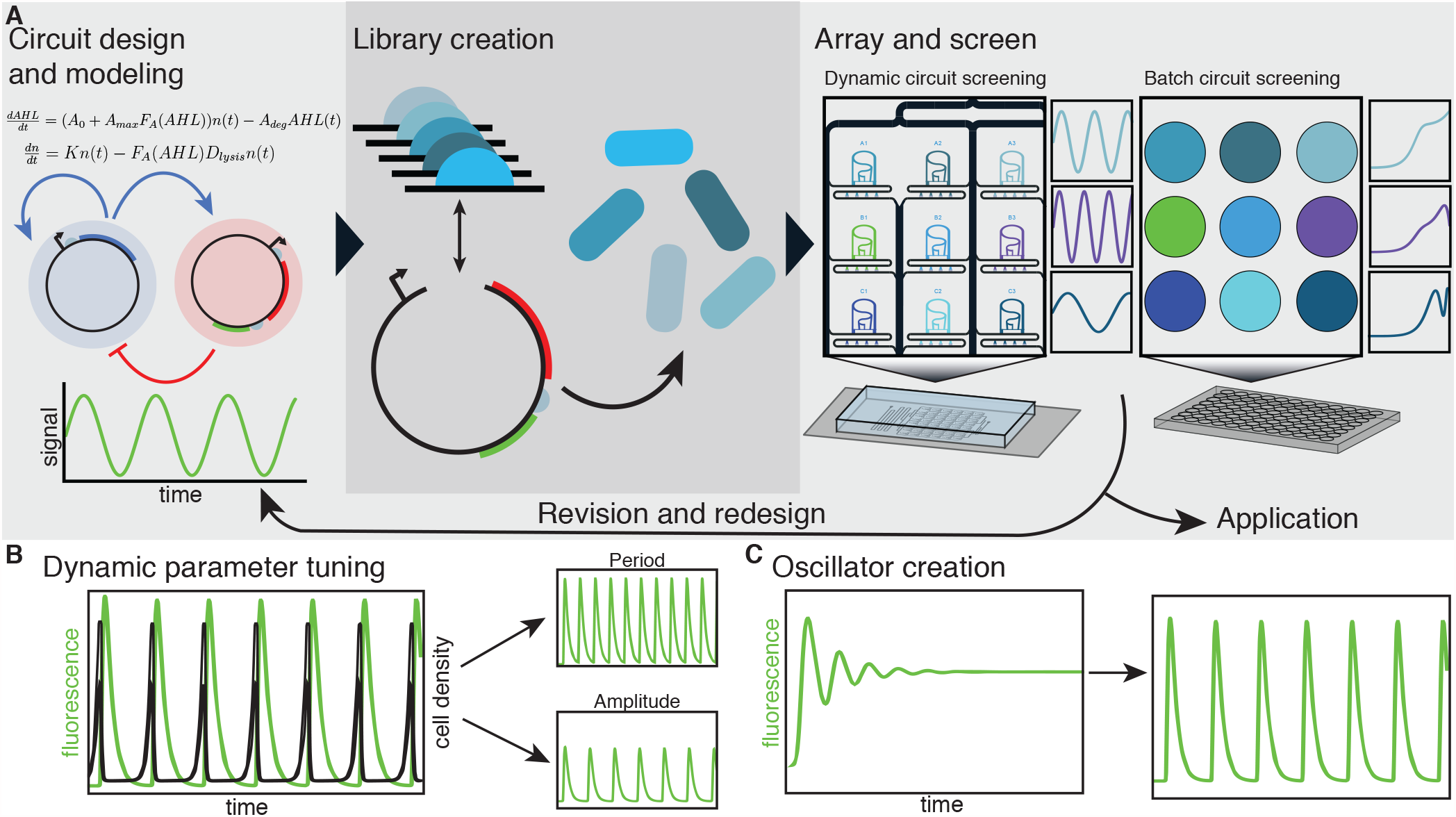
Synthetic oscillator creation and tuning through directed mutagenesis and screening. (A) Overview of gene circuit creation and screening work-flow. Mathematical modeling of circuit dynamics helps to identify parameters to target in order to improve or modify circuit behavior. Large libraries of a given circuit are quickly constructed via site-directed-mutagenesis (SDM). High-throughput, multi-strain microfluidic devices permit dynamic phenotype screening to supplement and improve upon traditional batch-culture methods of circuit screening. Circuit variants with desired or interesting behavior can be used for real-world applications, used to better inform circuit models, or placed through another cycle of mutagenesis and screening to further improve behavior. (B) The gene circuit library construction and screening work-flow developed here can be used to tune the behavior of an existing oscillator circuit. (C) The system can also aid in the construction of new genetic circuits such as oscillators synchronized at the population level.

### Tuning the oscillatory dynamics of a synchronized lysis circuit by directed mutagenesis

In order to demonstrate the ability of our system to tune dynamic gene circuits via directed mutagenesis and screening, we worked with a single plasmid version of a previously-developed synthetic gene oscillator, the synchronized lysis circuit (SLC)^25^. Bacteria transformed with the SLC have been used to release therapeutics in solid tumors^25–27^, and the ability to tune the circuit dynamics via directed mutagenesis could improve the utility of this circuit for cancer therapy. In the SLC, the expression of the LuxI protein, and subsequent production of the quorum sensing autoinducer N -Acyl homoserine lactone (AHL), generates synchronized positive feedback in a colony of isogenic cells. The positive activation of the pLuxI promoter in-turn drives the expression of negative feedback via the lysis protein, E from phage *ϕ*X174, causing the synchronized lysis of the colony. A few cells in the population are able to survive the synchronized lysis event and continue growing, perpetuating cycles of growth, gene expression, and mass lysis (Fig. 2A).

**Figure 2:**
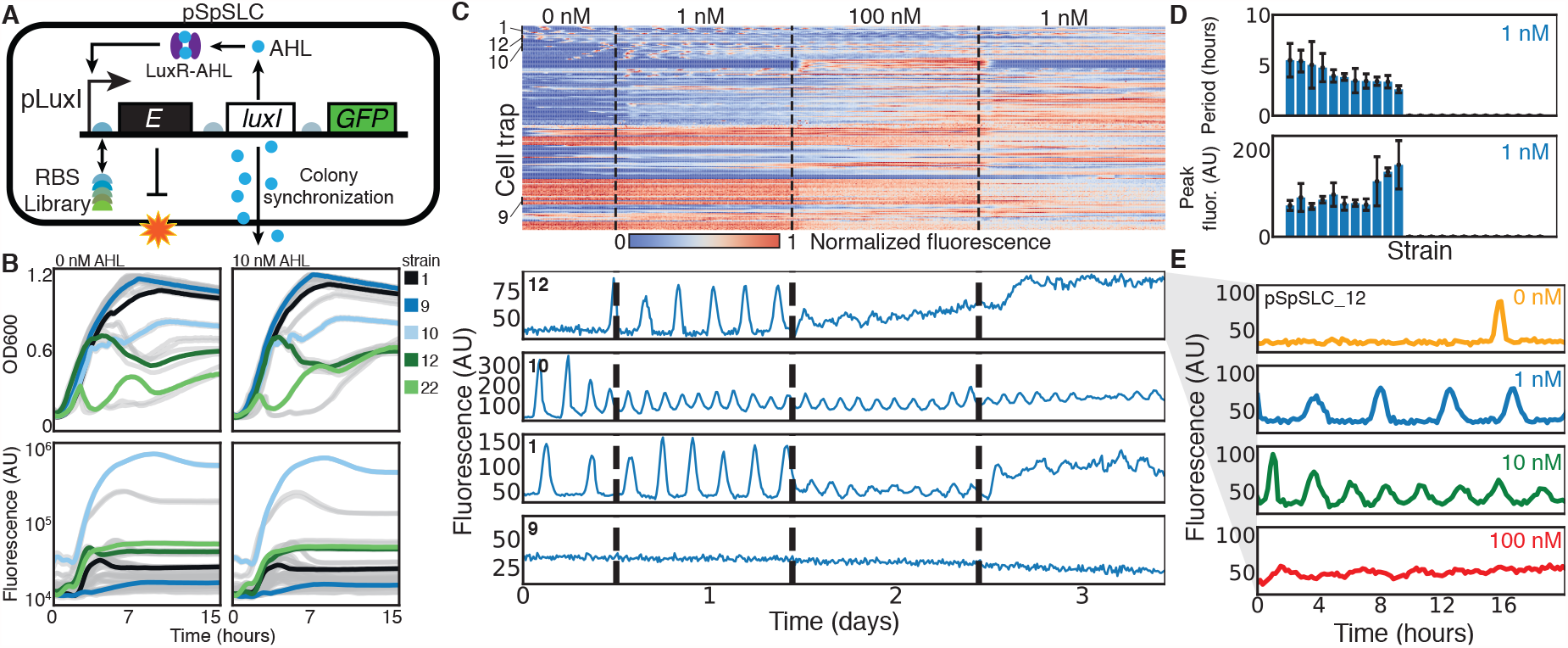
Screening of Synchronized Lysis Circuit (SLC) library strains. (A) A single-plasmid synthetic oscillator was developed with AHL production from the LuxI protein as a cell-synchronized positive feedback mechanism, and cell lysis as negative feedback. A library was created by randomizing five bases in the RBS upstream of the lysis protein, E. (B) When screened in batch culture, the library strains exhibit a range of growth, lysis and GFP expression dynamics. (C) 24 library members were screened on a 48 strain microfluidic device and subjected to temporal changes in the background AHL concentration. Different dynamic phenotypes were observed across these 24 strains, with four examples shown. (D) Extracted parameters of 24 oscillator strains under 1 nM AHL. The oscillatory period of each strain is shown, with 0 indicating no measured oscillations. The mean peak fluorescence values are shown below. Error bars represent standard deviation of measured cell traps. (E) A single strain subjected to multiple background concentrations of AHL exhibits varying dynamic phenotypes.

We generated a mutant library of the single plasmid SLC by randomizing five base pairs in the ribosome binding site (RBS) upstream of the lysis protein leading to as many as 1024 unique circuit variants (Fig. 2A). Altering the strength of the RBS preceding the lysis gene affects the translation rate for the lysis protein, which potentially alters the oscillatory dynamics of the circuit by modulating the negative feedback component of the circuit. We hypothesized that strains with a stronger RBS driving the lysis gene would lyse more rapidly upon reaching a threshold population size leading to higher frequency oscillations compared to a strain with a weaker RBS.

We randomly selected 24 members from the SLC library for screening. We cultured these strains in batch culture in a 96 well plate and monitored their lysis dynamics using a TECAN microplate reader. For the 24 strains examined in batch, we saw significant differences in the presence and magnitude of lysis events and GFP expression immediately before a lysis event (Fig. 2B). While differences in cell population dynamics and GFP fluorescence can be coarsely ascertained from the batch culture data, sustained SLC oscillations are typically only seen in continuous culture, necessitating the use of multiplexed microfluidics for dynamic parameter screening.

In parallel with the batch culture experiments, we screened the 24 selected library members on a 48-strain multiplexed device, with fluorescence measurements taken every 10 minutes. Over a period of 3 days, the device was subjected to varying background concentrations of AHL to determine its effect on oscillatory dynamics. Clustering of all cell traps reveals an abundance of phenotypes, predominantly “broken” oscillators with no oscillatory dynamics, but several working oscillators (Fig. 2C). Four strains are highlighted in Fig. 2C, showing oscillators that only activate under different AHL concentrations, dampen over time, or are continuously robust. Dynamic parameters (period and peak fluorescence) were extracted for all 24 strains at 1 nM AHL to demonstrate that a range of values could be generated and measured with this library and device (Fig. 2D). Finally, multiple chips were set up in parallel to observe the phenotype of these oscillator strains when grown under different AHL concentrations (Fig. 2C). We observe that for the highlighted strain, oscillations are sparse at 0 nM of AHL, regular at 1 nM and 10 nM with frequency increasing at higher concentrations, and absent at 100 nM (Fig. 2E). The trend of more frequent oscillations with increasing exogenous AHL concentration matched modeling results obtained using a deterministic model of the SLC (SI Fig. 1).

To better understand what was driving the dynamic differences between working oscillators, we investigated a subset in more detail. Specifically, we looked at the original oscillator we used to build the library which exhibits frequent oscillations with little to no GFP production before lysis and compared it to a library strain (#10) that exhibited slower oscillations and high GFP expression before each lysis event (Fig. 3A and B). To characterize how the different RBS’s for these two strains changed the lysis dynamics, we constructed strains where these RBS’s were cloned before the lysis gene in plasmids lacking the positive feedback component of the lysis circuit from the luxI gene (SI Fig. 9). We then characterized the lysis behavior of these strains in batch culture in response to varying AHL concentrations to generate lysis dose-response curves for the different RBS’s (Fig. 3C, D and SI Fig.4). We found that the original strain with more rapid oscillations had a much lower EC50 value of AHL to trigger lysis compared to the library strain with a longer period, demonstrating that this strain had a higher translation initiation rate for the lysis gene compared to library strain #10. Using a previously-developed deterministic model of the lysis circuit dynamics^28^, we confirmed that the period of oscillations is generally inversely correlated with the strength of expression for the lysis gene (Fig. 3 E,F, and G). Additionally, using this model, we found that the peak AHL (and GFP) production immediately preceding a lysis event decreased as the expression strength of the lysis gene was increased (Fig. 3 F and SI Fig. 2). This prediction from the model agreed with our experimental results where the library strain with the weaker RBS driving the lysis gene exhibited substantially more GFP expression preceding lysis.

**Figure 3:**
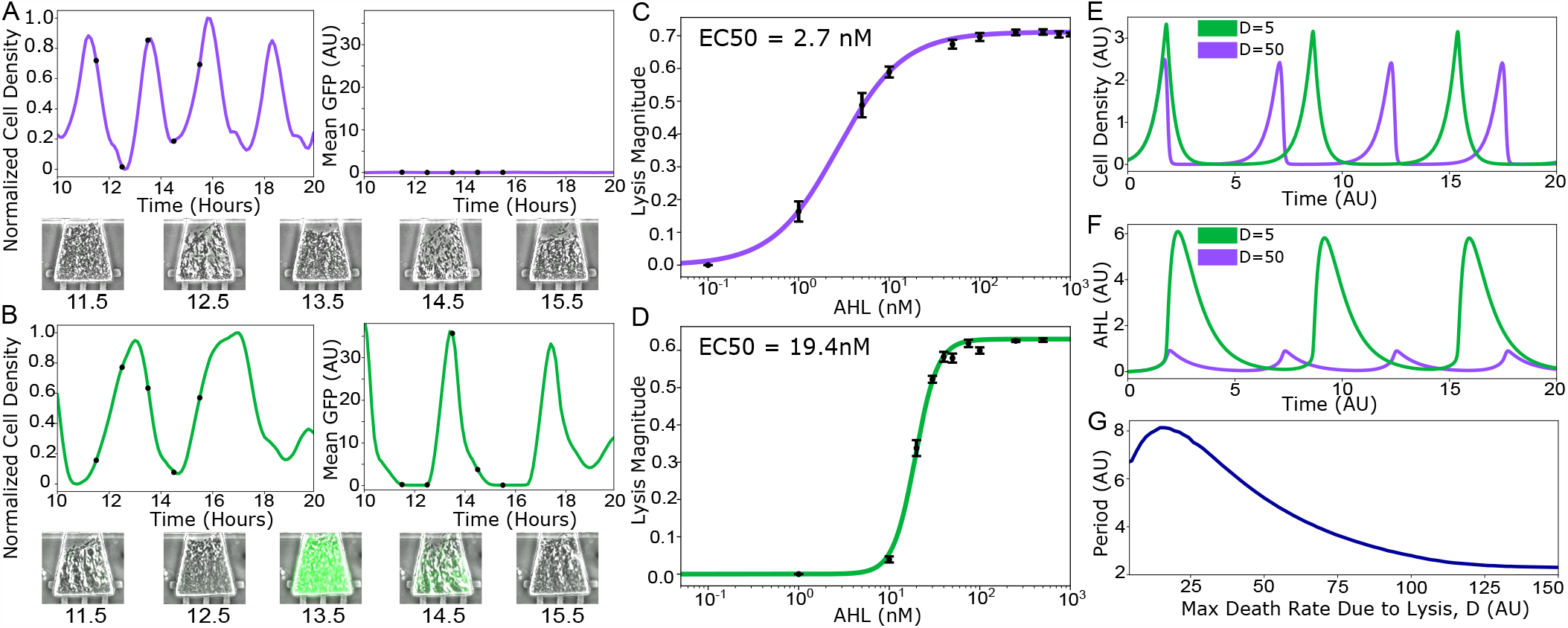
Comparison of SLC library strains with varying lysis strengths reveals interesting differences in gene expression dynamics. (A) GFP and cell density time traces for the original single plasmid SLC grown in the multistrain microfluidic device with accompanying microscope images for specific time points. (B) GFP and cell density time traces for SLC library strain 10 grown in the multi-strain microfluidic device with accompanying microscope images for specific time points. (C) Lysis dose-response curve for the original SLC strain. Error bars represent standard deviation of three separate lysis measurements. (D) Lysis dose-response curve for SLC library strain #10. Error bars represent standard deviation of three separate lysis measurements. (E) SLC modeling results showing how changing the maximum death rate due to lysis, D, impacts oscillatory population dynamics. (F) SLC modeling results showing how changing the maximum death rate due to lysis, D, impacts AHL concentration dynamics. (G) Modeling results showing how the period of lysis oscillations changes with the parameter D.

Our results here demonstrate the importance of considering the relative RBS strength for the lysis gene, particularly in cases where two genes are output by the oscillator circuit. In therapeutic applications using the SLC^25^, it may be desirable to have the production of a therapeutic gene driven by the same pLux promoter as the lysis gene. Here, we demonstrate the importance of considering the relative expression strength for the lysis gene and a therapy gene in this scenario. If the lysis gene translation initiation rate far exceeds that of the therapy gene, the engineered cells may exhibit robust cycles of growth and lysis without releasing a significant amount of therapeutic, akin to the case for the original SLC strain where there was little to no GFP production preceding each lysis event.

### Creation of a novel, robust synchronized gene oscillator circuit via directed mutagenesis and screening

To demonstrate the utility our circuit screening platform in the construction of new genetic circuits, we designed a novel synchronized oscillator which uses a transcriptional repressor as the negative feedback component rather than cellular lysis (Fig. 4A). The design of this new oscillator circuit uses two AHL inducible promoters: P1, activated by the LuxR-AHL complex and repressed by the tetracycline repressor protein, TetR, and P2, only activated by LuxR-AHL. In this new oscillator, P1 drives production of LuxI, which synthesizes AHL and drives positive feedback. P2 drives production of fluorescently-tagged TetR, which represses P1. As each cell accumulates high levels of TetR, the promoter P1 becomes inactivated leading to a steady decline in AHL as it’s removed from the population by fluid flow. Since AHL is now gone from the system, promoter P2 becomes inactive and TetR levels gradually decrease due to degradation and dilution and the cycle repeats.

**Figure 4:**
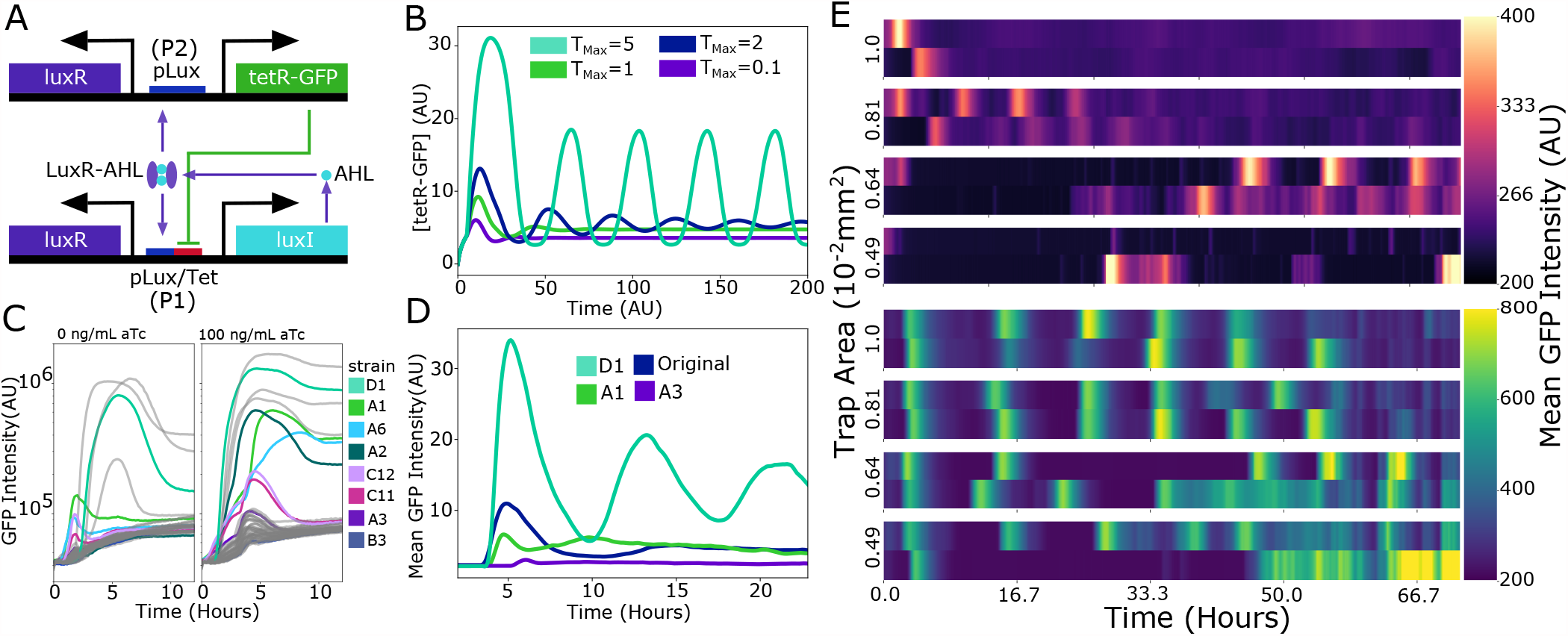
Creation of a new, robust synchronized gene oscillator via directed mutagenesis and screening. (A) A new synchronized oscillator circuit was designed based off of quorum sensing and transcriptional regulation. (B) Modeling results showing how the strength of TetR-GFP expression impacts oscillatory gene expression dynamics. (C) A library of potential oscillator strains was created by randomizing the RBS in front of the tetR-GFP gene by SDM. All of the strains were screened for differences in TetR-GFP expression in batch culture in the presence of 0 or 100ng/mL aTc. The eight highlighted strains were selected for additional characterization and testing in the multistrain microfluidic platform. (D) TetR-GFP time traces for a subset of strains screened in the multi-strain microfluidic platform. (E) Heatmaps showing oscillatory dynamics for the original implementation of the synchronized oscillator (top) and oscillator library strain D1 (bottom) in cell traps of different sizes.

We used a single-strain microfluidic device with variable cell trap sizes and a concentration gradient generator to characterize the dynamics of the first implementation of this oscillator design. The initial version of the circuit displayed oscillations in TetR-GFP expression with a period of about 7 hours in the microfluidic device, but these oscillations were relatively noisy across replicate cell traps (SI Fig. 5). Interestingly, in the largest microfluidic traps tested, the circuit exhibited one or two spikes in TetR-GFP expression before decaying to an intermediate level characterized by a mix of ‘ON’ and ‘OFF’ cells. To improve the consistency of oscillations, enable the circuit to function in larger populations and systematically investigate how changing circuit parameters affected the circuit, we made a small library of circuits where the strength of the ribosome binding site for TetR-GFP was varied. We chose to vary the strength of TetR expression based on computational modeling of the circuit suggesting that increasing TetR expression could move the circuit from a regime of damped oscillations to sustained oscillations (Fig. 4B).

To create a mutant library where the strength of TetR expression was varied, we changed the RBS preceding TetR to RBS sequences derived from the Anderson Lab RBS collection^29, 30^ by SDM. The final library consisted of as many as 4096 unique sequences. For initial screening of this circuit library, we picked 48 unique colonies and screened them in batch culture using a 96-well plate in the presence and absence of 100 ng/mL anhydrotetracycline (aTc) and tracked their GFP expression during growth (Fig. 4C). For further screening in microfluidics, we selected 8 library members that spanned the range of GFP expression we saw in the well plate assay and loaded these strains on the multi-strain microfluidic platform. When grown in the hydro-dynamically loaded cell traps of the microfluidic device, the majority of library strains exhibited one or two small peaks in TetR-GFP expression before decaying to relatively steady, intermediate levels of expression (Fig. 4D). One strain, D1, that had the strongest RBS driving TetR-GFP expression, was closer to having sustained oscillations in this microfluidic platform so we chose this strain to investigate further in a single strain microfluidic device.

When strain D1 was grown in the microfluidic chip with variable trap sizes, we found that it exhibited robust oscillations over long time periods in multiple trap sizes (Fig. 4E). Additionally, we found that the characteristics of the oscillations were relatively unaffected by aTc concentrations ranging from 0 to 50 ng/mL (SI Fig. 6). The period of the D1 oscillator was also able to be tuned significantly by varying the flow rate, with reduced flow rate leading to more frequent oscillations (SI Fig. 3). Lastly, when this new D1 oscillator strain was compared to the initial synchronized oscillator strain, we saw that the amplitude of oscillations for the new strain was greater and that the oscillations for the new strain were less noisy across replicate cell traps compared to the original strain (Fig. 4E, SI Fig. 5, and SI Fig. 6).

## Discussion

Tuning genetic circuit outputs by screening variant libraries for a desired phenotype has long been fundamental to synthetic biology design. However, the mass-screening of dynamic phenotypes has remained a persistent challenge and our ability to generate genotypic diversity far exceeds our ability to screen complex phenotypes^22^. Despite limited means for dynamic phenotype screening, canonical gene circuit motifs, including oscillators, logic gates, and feedback controllers have been increasingly deployed in time-dependent applications spanning metabolic engineering to therapeutic delivery ^25, 31–34^. Multiplexed microfluidics, such as ours, can aid in the development of circuits like these, for both academic research and as synthetic biology exits the lab and enters the real world.

For bacteria, it is well-documented that the growth-state of a growing culture has a significant impact on gene expression^35–37^. Thus, to reliably characterize and evaluate complex circuit dynamics, the cellular growth environment should be as constant as possible. In this article, we further demonstrate the importance of continuous culture screening, specifically in the context of dynamic gene circuits like oscillators. In screening the SLC library, we saw that the presence of a lysis event in the batch culture screen generally correlated with a propensity for robust oscillations in continuous culture, but continuous culture was necessary to confirm and detect sustained oscillations for any library members. On the other hand, batch screening of the TetR-GFP synchronized oscillator provided little evidence regarding which strains were more likely to oscillate in microfluidic culture but did facilitate the selection of strains with varying TetR expression for further testing in microfluidics. While our results show that batch culture can offer some insight into the design of oscillator circuits, in this context, batch culture is most useful as a means to cull non-interesting variation in dynamic gene circuit libraries. Ultimately, the microfluidic approach is necessary for fully characterizing dynamic phenotypes.

Microfluidics has served as useful tool for approximating complex real world environments in the past, simulating environments spanning soil to human organs^38, 39^. While not a perfect recreation of these complex environments, tuning environmental and time-dependent parameters achievable with microfluidics serves an important role in the prototyping and scale-up of all classes of gene circuits. The SLC library presented in this study, as an example, demonstrated variance between the magnitude of lysis events and the expression of a reporter gene. This could play a key role in its therapeutic deployment^25^, where relative dynamics of lysis and the expression of a therapy gene are critical to circuit success. Furthermore, this work shows how environment, specifically population size, can significantly impact the circuit dynamics, with the TetR synchronized oscillator behaving differently when grown in different cell trap geometries. Understanding how circuit dynamics change, or are resistant to change, as trap size varies can be critical to predicting how a circuit might behave when deployed in a real world, non-microfluidic environment.

It is well-documented that small changes to the RBS preceding a gene can cause dramatic impacts on gene expression^40^, and the small size of many RBS sequences makes them an easy target for SDM. Through creating and screening mutant oscillator libraries, we saw that modifying a single RBS sequence in the circuit by SDM can dramatically change circuit dynamics. For the SLC, changing the RBS before the lysis gene was able to change the period of oscillations and the circuit’s sensitivity to exogenous AHL. For the TetR-GFP synchronized oscillator, changing the RBS before the *tetR* gene moved the circuit dynamics from damped to sustained oscillations. Although we only targeted RBS sequences for library creation in this work, other circuit components could easily be targeted with similar methods such as promoter sequences, operator sites, and coding DNA sequences. Additionally, the multiplexed microfluidic technology we used would be easily amenable to screening genetic circuits in different host strains of *E. coli* and potentially other species of bacteria as well.

The final version of the TetR-GFP synchronized oscillator provides a simple blueprint for building future oscillators. Previous synchronized oscillators developed in our laboratory have all utilized autoinduction of an AHL synthase gene for positive feedback and synchronization, but have used a range of sources for negative feedback including: 1) enzymatic degradation of AHL^41^, 2) restriction enzyme cutting of plasmid DNA^42^, and 3) cell lysis^25^. Here we show that a synthetic oscillator design based off of quorum sensing and simple transcriptional regulation can be used to generate synchronized oscillations. Presumably this circuit architecture could be easily adapted to different quorum sensing systems, expanding its utility and paving the way for synchronized oscillator circuits in more diverse species of bacteria.

## Methods

### Microfluidic device development and fabrication

Our group has previously described the microfabrication techniques used to pattern SU-8 photoresist onto a silicon wafer to create the mold for our device ^43^. A poly-dimethylsiloxane (PDMS) device was made from the wafer by mixing 77 grams of Sylgard 184 and pouring it on the wafer centered on a level 5”x5” glass plate surrounded with an aluminum foil seal. The degassed wafer and PDMS was cured on a flat surface for one hour at 95°C.

### Multi-strain microfluidic device loading and bonding

A PDMS device cleaned with 70% Ethanol and adhesive tape was aligned to a custom fixture compatible with the Labcyte Echo. Both the fixture and a clean glass slide sonicated with 2% Helmanex III were exposed to oxygen plasma. *E. coli* to be used in microfluidic experiments were grown for 16 hours on LB media, after which 45 *µ*L of each strain was pipetted into a well of a Labcyte Echo 384-well microplate and 2.5 nL was acoustically transferred to each device position on the PDMS device. The spotted device and glass slide were bonded together and cured at 37°C for two hours.

### Single strain microfluidic device loading and bonding

For the single-strain microfluidic experiments (Fig.4E), a previously-developed PDMS device with variable cell trap sizes and a concentration gradient generator was used ^44^. Prior to cell loading, the device was placed in a vacuum chamber for 30 minutes. During this period, 1mL of an overnight culture of the engineered strain was spun down and concentrated in 10*µ*L of LB media with 0.075% tween. Immediately following removal from the chamber, the cell suspension was pipetted to cover the outlet of the device and sterile LB media + 0.075% tween was pippetted to cover the two inlet ports. After media and cells were pulled into the microfluidic chip by the vacuum and the cell traps had filled with cells, two inlet syringes with fluidic tubing attached were connected to the inlet ports of the device. Similarly an outlet syringe with tubing was connected to the outlet port of the device. All of the cell traps had the same width (100*µ*m) and height (1.2*µ*m) and ranged in length from 40 to 100*µ*m. Media flow was maintained across the device by maintaing the source syringes 5-10 inches above the outlet syringe fluid height. For experiments using anhydrotetracycline (aTc), one inlet syringe was prepared with a concentration of 50ng/mL aTc in LB while the other syringe was prepared with 0ng/mL aTc leading to a gradient of 8 different aTc concentrations across the device.

### Microfluidic experimental protocol

Microfludic experiments were performed on a custom optical enclosure described in the supplemental information or on a Nikon TE2000-U epifluorescent inverted microscope (Nikon Instruments Inc., Tokyo, Japan). Cells were grown on the device on LB media with the appropriate antibiotics, and 0.075% Tween-20 until traps were filled to confluence. Extracted fluorescence time series were normalized to remove device background fluorescence and strain background fluorescence. Detailed methods on experimental set-up and data collection can be found in the supplement.

### Cloning and mutant library generation

The original versions of the synchronized oscillator strains were cloned using Gibson assembly. Plasmid sequences were confirmed with Sanger sequencing (Eton Bioscience, San Diego, CA).

To generate a mutant library of the single plasmid SLC oscillator plasmid (pSpSLC), 5 base pairs in the Shine-Dalgarno sequence of the ribosome binding site (originally GAGAA, located 7 to 12 base pairs upstream of the start codon of the lysis protein, E) were randomized by site directed mutagenesis. The entire plasmid was PCR amplified using Q5 DNA Polymerase (New England BioLabs Inc.) with the following degenerate primers where N indicates any base: 5’ CATTAAAGAGNNNNNAGGTACCATGATGGTAC 3’ and 5’ AATTCTCTCTATCACTGATAG 3’. Degenerate primers were ordered from Integrated DNA Technologies (IDT). A blunt-end ligation was performed to re-circularize the plasmid before transformation into MG1655 *E. coli* competent cells. 24 colonies from the agar plate were randomly selected for mutant screening and grown up for 16 hours in LB media with 0.2% glucose and spectinomycin prior to use in experiments.

To generate a mutant library of the two plasmid TetR-GFP synchronized oscillator, the RBS preceding TetR-GFP on the plasmid pTetSO2 was randomized to RBS sequences derived from an RBS library created by Professor Christopher Anderson^29, 30^. Specifically, the entire plasmid was PCR amplified using Q5 DNA Polymerase with the following degenerate primers where N indicates any base: 5’ NNGANNNACTAGATGTCTAGATTAGATAAAAGTAAAG 3’ and 5’ NTCTTTCTCTAGAATTCGACTATAACAAACCATTTTC3’. Degenerate primers were ordered from Integrated DNA Technologies (IDT). A blunt-end ligation was performed to re-circularize the plasmid before co-transformation with plasmid pTetSO1 into MG1655 *E. coli* competent cells. The transformation was plated on LB agar containing chloramphenicol and spectinomycin and 48 colonies from the agar plate were randomly selected for mutant screening.

### Generation of lysis dose-response curves

To generate the lysis dose-response curves shown in Figure 3, 200*µ*L cultures of the strains containing the plasmid pAHL Lyse were started by seeding cells from a saturated culture at a 1:100 ratio in LB media. The cultures were grown at 37C and optical density was monitored every five minutes using a TECAN microplate reader with orbital shaking. Once the cultures reached early exponential phase (OD 0.2-0.3), the well plate was quickly removed them the microplate reader and each culture well was spiked with 2*µ*L of a 100X AHL stock to achieve the desired final concentration. The well plate was then re-inserted into the microplate reader and the cultures were grown for 12 hours.

To generate a lysis magnitude value from each condition, the growth curve for that condition was examined for an inflection point where the derivative of the culture OD with respect to time changed from positive to negative. Then, lysis magnitude was calculated as L/G, where G is the positive change in OD from the initial time point to the inflection point and L is the negative change in OD from the inflection point to the time point at which OD was lowest following the inflection point (SI Fig. 4). A dose response curve was generated with the lysis magnitude vs. AHL concentration data by fitting the data to the Hill Equation.

### Data analysis of multi-strain microfluidic TL stacks

To calculate the normalized cell density vs. time plots shown in Figure 3 using 10x transmitted light (TL) microscope images, the following protocol was used. First, the mean TL pixel value for each trap (*TL*_*trap*_) was extracted in ImageJ along with a the mean pixel value for a selection of equal size on a part of the chip containing no cells (*TL*_*BG*_). To obtain an approximate cell density from these two measurements, the following formula was used: 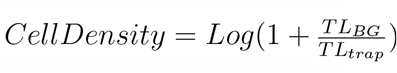. Lastly, for each trap the approximate cell density was normalized as: 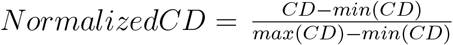

### Analysis of single strain microfluidic data

To analyze the 10x image stacks obtained from time-lapse microscopy experiments with the TetR-GFP synchronized oscillator strains, the mean GFP pixel intensity from each image was extracted using imageJ for each trap size and inducer concentration. The data shown in SI Fig. 5 and 6 represent the mean GFP data for two separate microfluidic experiments. To calculate the average oscillatory period for the D1 oscillator for different conditions (SI Fig. 3), cell traps were chosen that had at least two distinct peaks in mean GFP expression for each flow rate and trap size combination. The period for each cell trap that was included in the data analysis was calculated as the mean time elapsed between each peak divided by the number of GFP peaks. The bar plot in SI Fig. 3 was created by taking the mean period for each flow rate and trap combination and the error bars represent the standard deviation among analyzed cell traps.

### Deterministic modeling of Synchronized Lysis Circuit dynamics

For all modeling of the SLC, we used a modified version of a previously-published deterministic model of SLC dynamics ^28^. This simple model consists of two differential equations, one that describes the production and dilution of the quorum sensing molecule AHL (Equation 1) and one that describes cell growth and lysis-induced cell death (Equation 2). We added an additional ODE to this model to directly account for AHL-induced GFP production (Equation 3). To model the effect of exogeneous AHL on circuit dynamics (SI Fig. 1), we modified equation 1 so that the value of AHL at any given time-point (*AHL*(*t*)) was not allowed to decrease below some set point *AHL*_*min*_. All model results were obtained in MATLAB using the ode45 function. The following parameters were used in the SLC model simulations except where noted: *K* = 2, *D*_*lysis*_ = 5, *A*_0_ = 0.4, *A*_*max*_ = 8, *AHL*_*th*_ = 1, *m* = 4, *A*_*deg*_ = 1, *G*_*max*_ = 8, *G*_*deg*_ = 1, *GFP*_*th*_ = 3

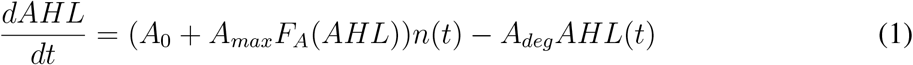

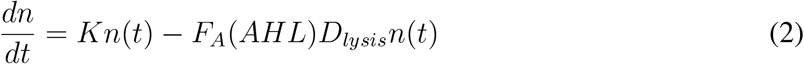

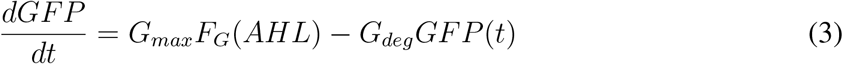

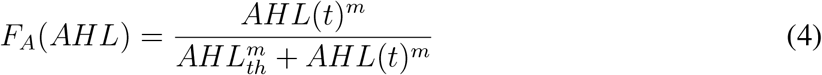

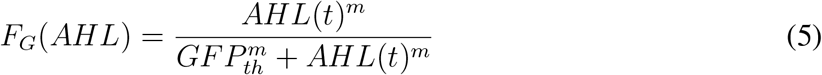

### Deterministic modeling of tetR-GFP Synchronized Oscillator dynamics

To model the behavior of the TetR-GFP synchronized oscillator, we used a delayed, ordinary differential equation model loosely based off a previous model of a similar synchronized oscillator^41^. The model consists of two main equations describing the production and degradation of AHL (Equation 6) and tetR (Equation 7). Equation 6 takes into account that both the basal and maximal production rates of AHL are affected by TetR repression while basal AHL production leads to additional AHL production in an auto-catalytic positive feedback loop. Equation 7 takes into account that TetR expression is only impacted by AHL. In the model, the degradation terms for both AHL and TetR represent that both proteins are actively degraded by the same protease (ClpXP) via Michaelis-Menten kinetics. To account for delays in the transcription, translation, and production of AHL and TetR relative to their rapid binding to transcription factors or operator sites, we include a delay term (*τ*) in the hill functions for AHL and TetR (Equations 9, 8). All model results were obtained in MATLAB using the delayed differential equation solver, solveDDE. The following parameters were used in all of the synchronized oscillator model simulations except where noted: *A*_0_ = 5, *T*_0_ = 2, *A*_*max*_ = 30, *T*_*M*_ *ax* = 3, *AHL*_*th*_ = 1, *tetR*_*th*_ = 1, *m* = 2, *A*_*deg*_ = 1, *T*_*deg*_ = 1, *f*_*deg*_ = 0.1, *τ* = 5

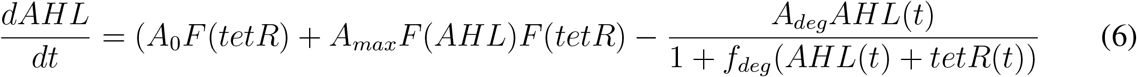

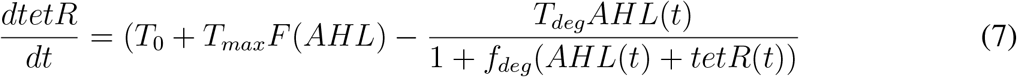

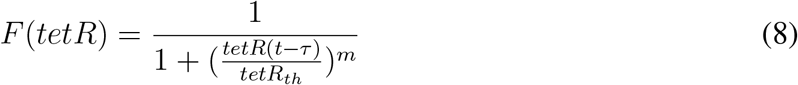

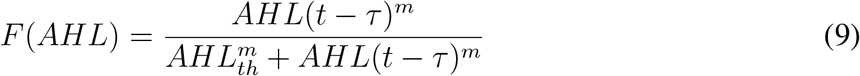

## Abbreviations

SLC: Synchronized Lysis Circuit
RBS: Ribosome Binding Site
SDM: Site Directed Mutagenesis

## Acknowledgements

This work was supported by the Defense Advanced Research Projects Agency (DARPA). The authors thank Arianna Miano for providing the device for single strain microfluidic experiments, Jaquelin Dezha Peralta for assistance with microfluidic fabrication, and Shalni Kumar for critical reading of the manuscript.

## Author Contributions

Author contributions: Conceptualization, A.L., N.C., and J.H.; Investigation, A.L. and N.C.; Writing and Visualization, A.L., N.C., and J.H.

## SI Materials and methods

### Microfluidic device fabrication

A poly-dimethylsiloxane (PDMS) device was made from the wafer by mixing 77 grams of Sylgard 184 and pouring it on the wafer centered on a level 5”x5” glass plate surrounded with an aluminum foil seal. The degassed wafer and PDMS was cured on a flat surface for one hour at 95°C.

### Cloning and mutant library generation

To generate a mutant library of the pSpSLC oscillator plasmid, 5 base pairs in the Shine-Dalgarno sequence of the ribosome binding site (located 7 to 12 base pairs upstream of the start codon of the lysis protein, E) were randomized by site directed mutagenesis. The original sequence at this position was: GAGAA. First, the entire plasmid was PCR amplified with the following degenerate primers where N indicates any base: 5’ CATTAAAGAGNNNNNAGGTACCATGATG-GTAC 3’ and 5’ AATTCTCTCTATCACTGATAG 3’. The PCR reaction mix was incubated with DPNI at 37C for 30 minutes to digest template plasmid and then the 4.7 kb PCR product was run on an agarose gel and extracted using a QIAquick Gel Extraction Kit (QIAGEN). 1*µ*L of the gel extracted PCR product was mixed with 0.5*µ*L T4 ligase buffer, 0.5*µ*L T4 PNK, and 3*µ*L of DNase-free water and incubated at 37C. Next, 0.5*µ*L T4 ligase buffer, 0.5*µ*L T4 DNA ligase, and 4*mu*L were added to the reaction mixture and the mixture was incubated at room temperature overnight. The following day, 50*µ*L of chemically competent MG1655 E. coli cells were transformed with 3*µ*L of the reaction mix and plated on an LB agar plate containing 0.2% glucose and spectinomycin. 24 colonies from the agar plate were randomly selected for mutant screening and grown up for 16 hours in LB media with 0.2% glucose and spectinomycin prior to use in experiments.

### Cell preparation

For multi-strain microfluidic experiments, cells were grown overnight on LB+antibiotic media. Lysis oscillator strains were grown on LB supplemented with 0.2% glucose to suppress expression of the pLux promoter driving lysis. Turbid cultures were transferred to a 384 Echo compatible plate for direct transfer onto microfluidic devices.

### Microfluidic device loading and bonding

A PDMS device cleaned with 70% Ethanol and adhesive tape was aligned to a custom fixture compatible with the Labcyte Echo. Both the fixture and a clean 4”x3” glass slide sonicated with 2% Helmanex III were exposed to oxygen plasma. Cells were spotted from the 384 Echo compatible plate directly onto the PDMS devices. The device and glass slide were bonded together and cured at 37°C for two hours.

### Multi-strain microfluidic device experimental set-up

Before setting up a microfluidic experiment, the device was placed in a vacuum for a minimum of 20 minutes. The device was then mounted onto the custom optical enclosure. The inlet port was connected to a 50 mL syringe and tygon tubing with LB media with antibiotic (spectinomycin for SLC oscillator strains, chloramphenicol and spectinomycin for tetR-GFP synchronized oscillator strains), and 0.075% Tween-20. The waste port was connected to tygon tubing and a 1L waste bottle. The height difference between the inlet and outlet was 20” corresponding to a flow rate of approximately 1 mL/hr. Tween-20 was used in the media as a surfactant to reduce clogging and therefore increase the longevity of microfluidic experiments. Tween-20 has been used by our group in many experiments without an adverse effect on *E. coli* ^43, 45^.

### Induction protocol

For AHL inductions, LB with the predetermined AHL concentration was mixed and pipetted into the source media syringe. For periods where the same AHL concentration was left on the device for over 24 hours, the media was pipetted out of the syringe and replaced.

### Live-cell imaging and data extraction

Microfluidic devices were imaged in a custom optical enclosure continuously every ten minutes in both the transmitted light and gfp fluorescence channels with a 1 second and 60 second exposure respectively. The custom optical enclosure uses an SBIG STX-16803 CCD Camera with a custom lens stack assembly containing the Semrock FF01-466/40-32-D-EB and FF02-520/28-50-D-EB excitation and emission filters, respectively. The enclosure has green and blue LED spotlight sources obtained from ProPhotonix for transmitted light and fluorescence light sources, respectively. The optical resolution of the enclosure is 36 *µ*m. The enclosure was temperature controlled to 37°C.

Images were synced from the enclosure to a server via WiFi for further data processing. Custom software produced flat-field-corrected images in both channels in real-time to remove optical vignetting using the following equation:

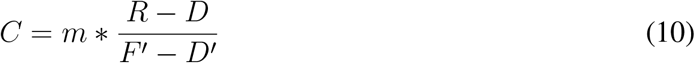

where *R* is the raw image to be flat-field corrected, *D* is the dark-current image for that device, taken at the same exposure settings as *R, F* is a raw image taken by the camera with no device present, *D*′ is the dark-current image taken at same exposure as *F*′, *m* is the mean value for all values in the array (*F′ − D′*), and *C* is the resulting corrected image.

Flat-field corrected images were then processed in ImageJ, where a custom ”Region Of Interest” or ROI manager was used to extract fluorescence, transmitted light, and background values.

Data was initially processed by subtracting the local background signal, in order to eliminate any local or regional fluctuations that are of an additive (or, analogously, subtractive) nature. The result of this background correction was to produce a vector 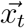 representing the background-corrected fluorescent signals of all cell traps at time *t*:

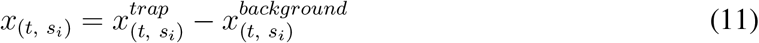

where *t* refers to the current time point, *s*_*i*_ refers to the strain in cell trap 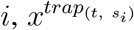 is the flat-field corrected fluorescent signal from the *trap* of position *i* at time *t*, and 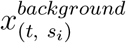 refers to the flat-field corrected local background fluorescent signal at position *i* at time *t*.

## SI Figures

**SI Figure 1:**
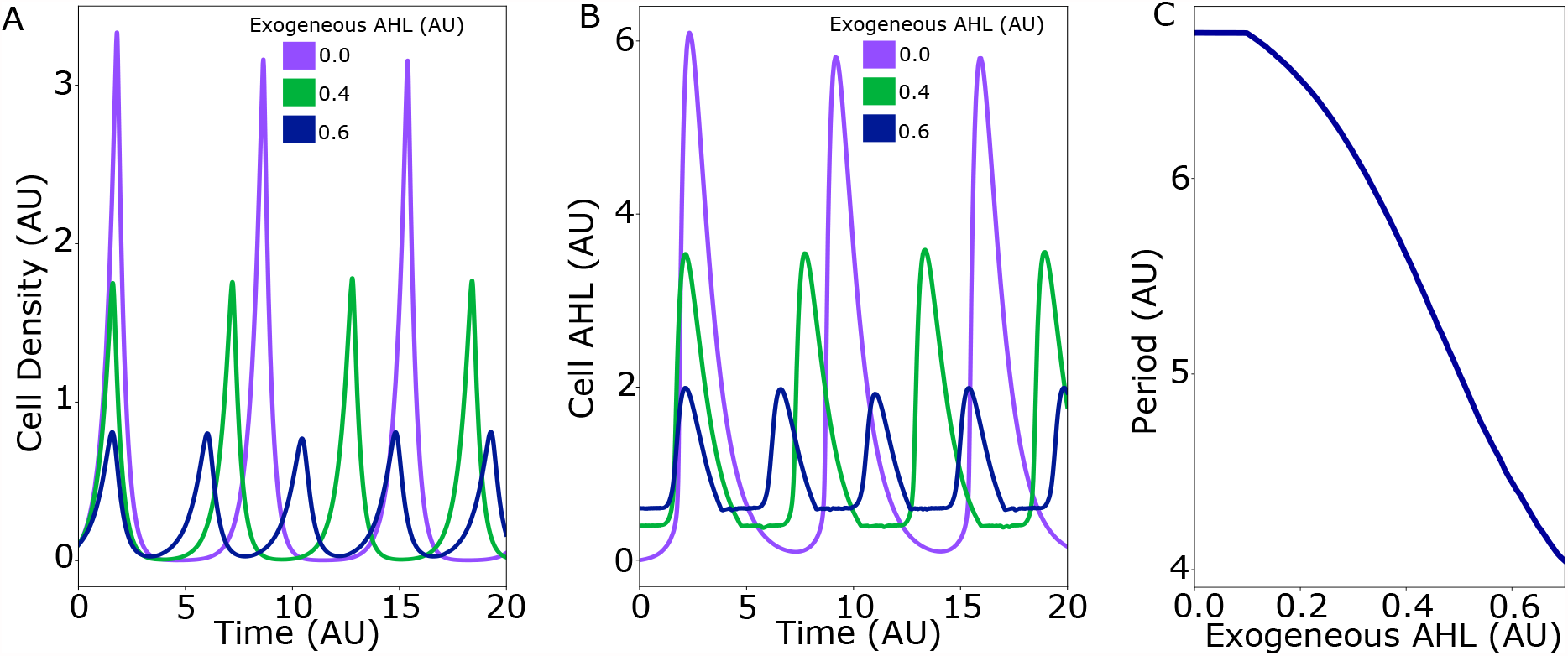
Modeling the effect of exogeneous AHL on SLC oscillatory dynamics. (A) Cell density vs. time traces for three different exogenous AHL values. (B) Cellular AHL concentration vs. time traces for three different exogenous AHL values. (C) Oscillatory period vs. Exogeneous AHL concentration

**SI Figure 2:**
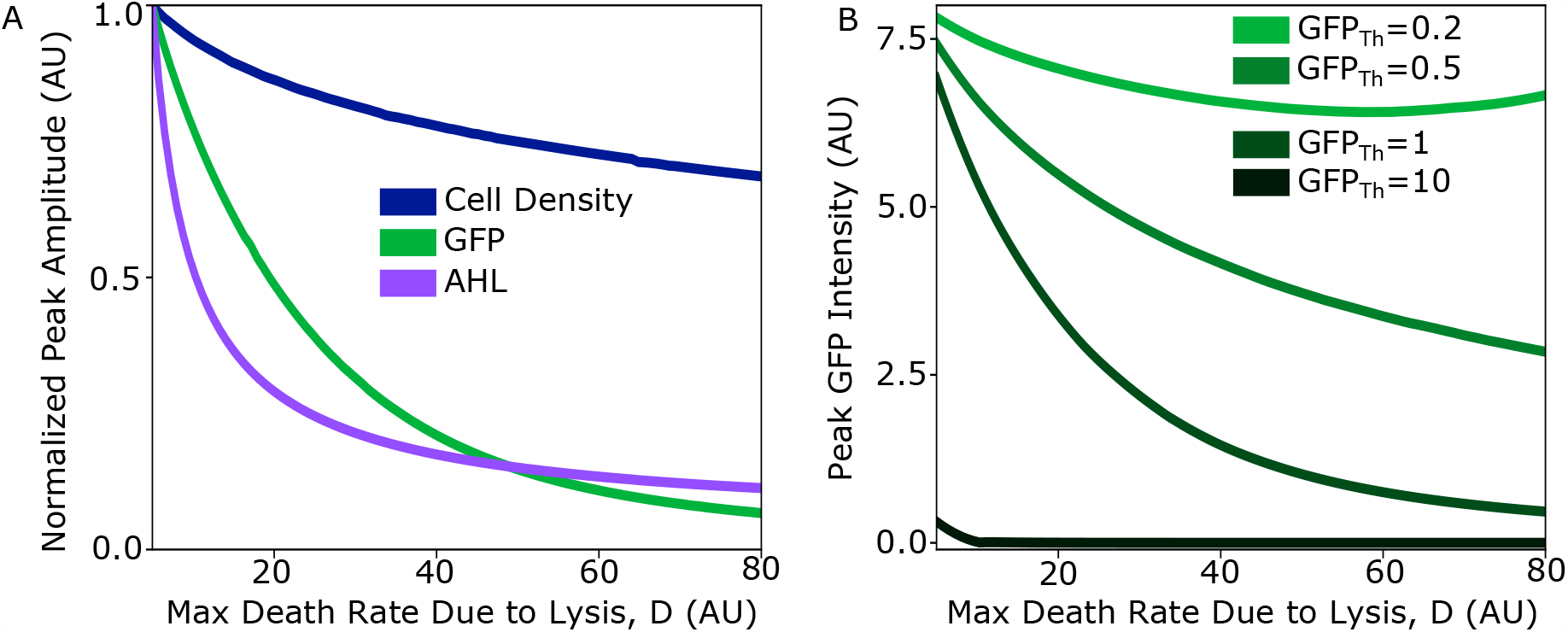
Modeling the effect of lysis strength on GFP expresion in the SLC. (A) Normalized peak amplitude of cell density, cellular AHL, and cellular GFP for varying values of the model parameter *D*, the max death rate due to lysis. (B) Peak GFP intensity as a function of D for varying threshold AHL values needed to trigger GFP expression, *GFP*_*Th*_

**SI Figure 3:**
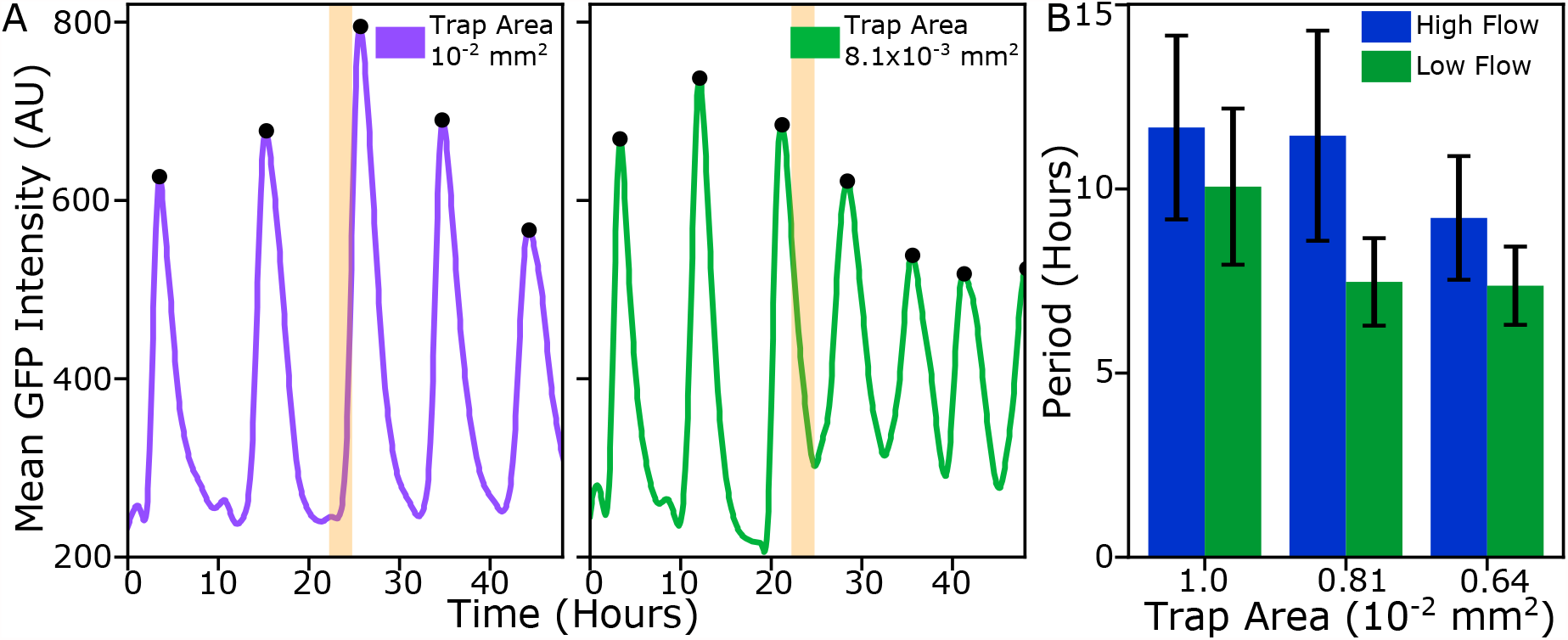
The improved TetR-GFP synchronized oscillator oscillates with different frequencies based on trap size and flow rate (A) Representative time traces of oscillator GFP dynamics in two different cell trap sizes. Light red vertical bars indicate time point where the hydrostatic pressure driving flow was reduced from 10 to 6 inches of *H*_2_*O* (B) Mean period of oscillations for three different trap sizes at high and low flow rates corresponding to hydrostatic pressure of 10 and 6 inches of *H*_2_*O* respectively

**SI Figure 4:**
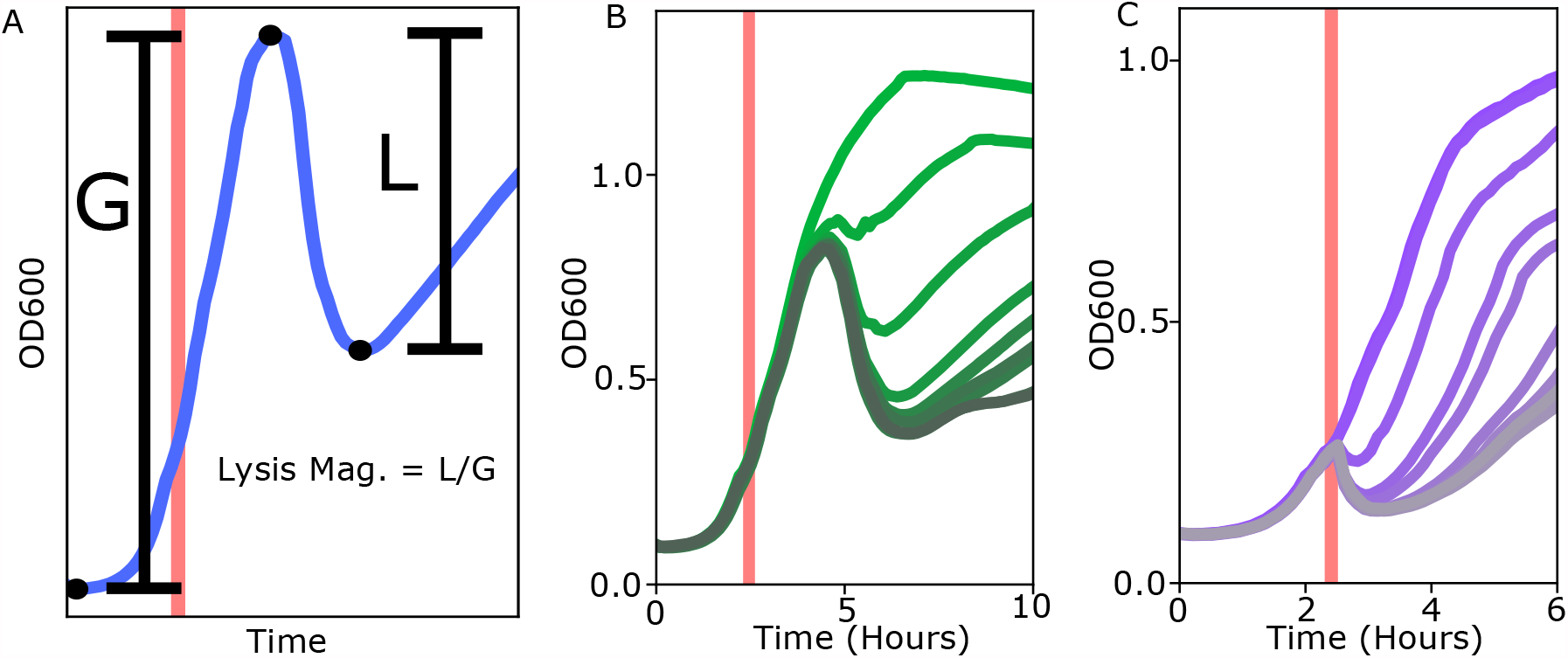
Method for creating lysis dose-response curves to compare expression strength of different RBS sequences driving lysis (A) Lysis magnitude was calculated as the change in culture optical density (OD) during a lysis phase (L) divided by the change in OD during a growth phase (G). Light red vertical bar represents time point when 2uL of a 100X AHL stock was spiked into each culture well to achieve the desired AHL concentration. (B) Representative growth and lysis curves used to construct the dose-response curves shown in Figure 3. Green traces represent SLC library strain #10 while purple traces are for the original SLC strain.

**SI Figure 5:**
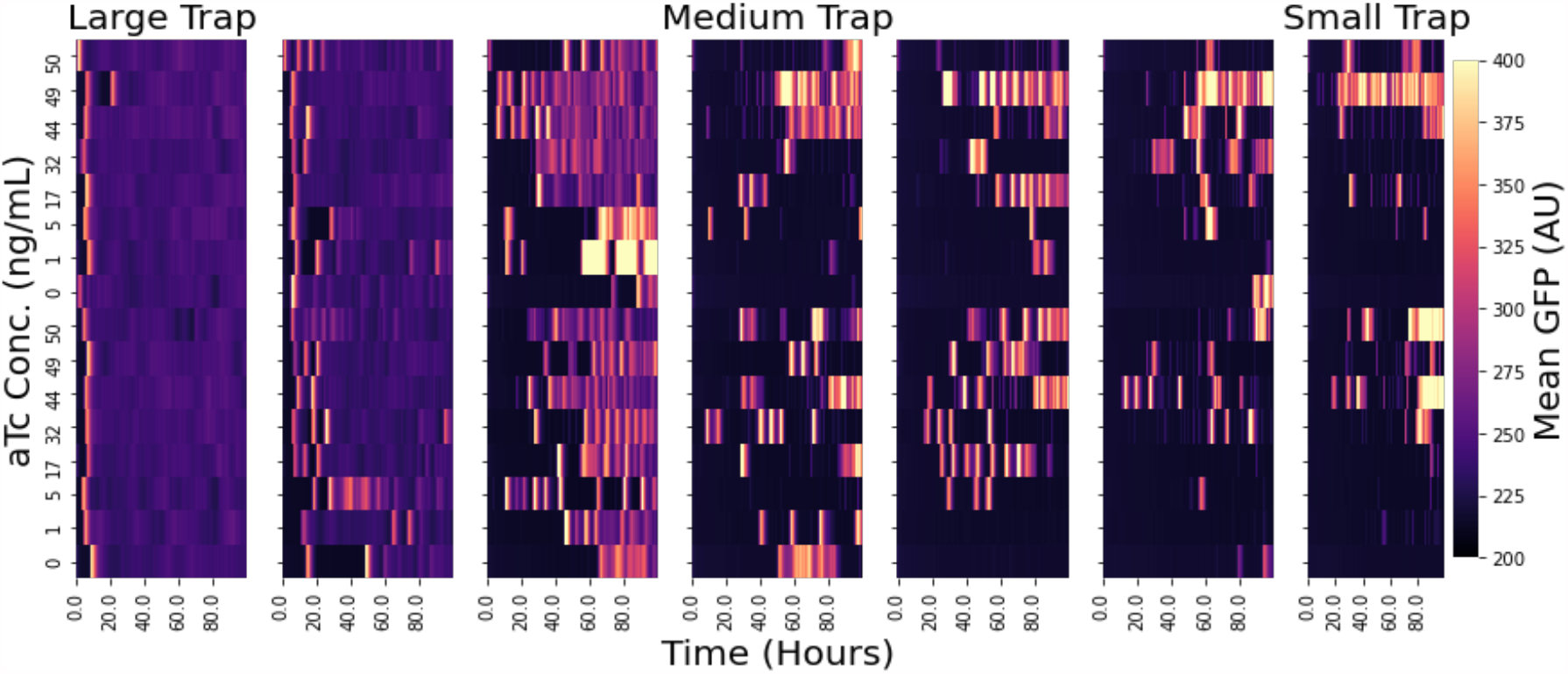
Full single strain microfluidic data set for the original implementation of the TetR-GFP synchronized oscillator. Cell trap sizes ranged from 1.0 *\** 10^*−*2^*mm*^2^ (Large Trap) to 0.16 *\** 10^*−*2^*mm*^2^ (Small Trap)

**SI Figure 6:**
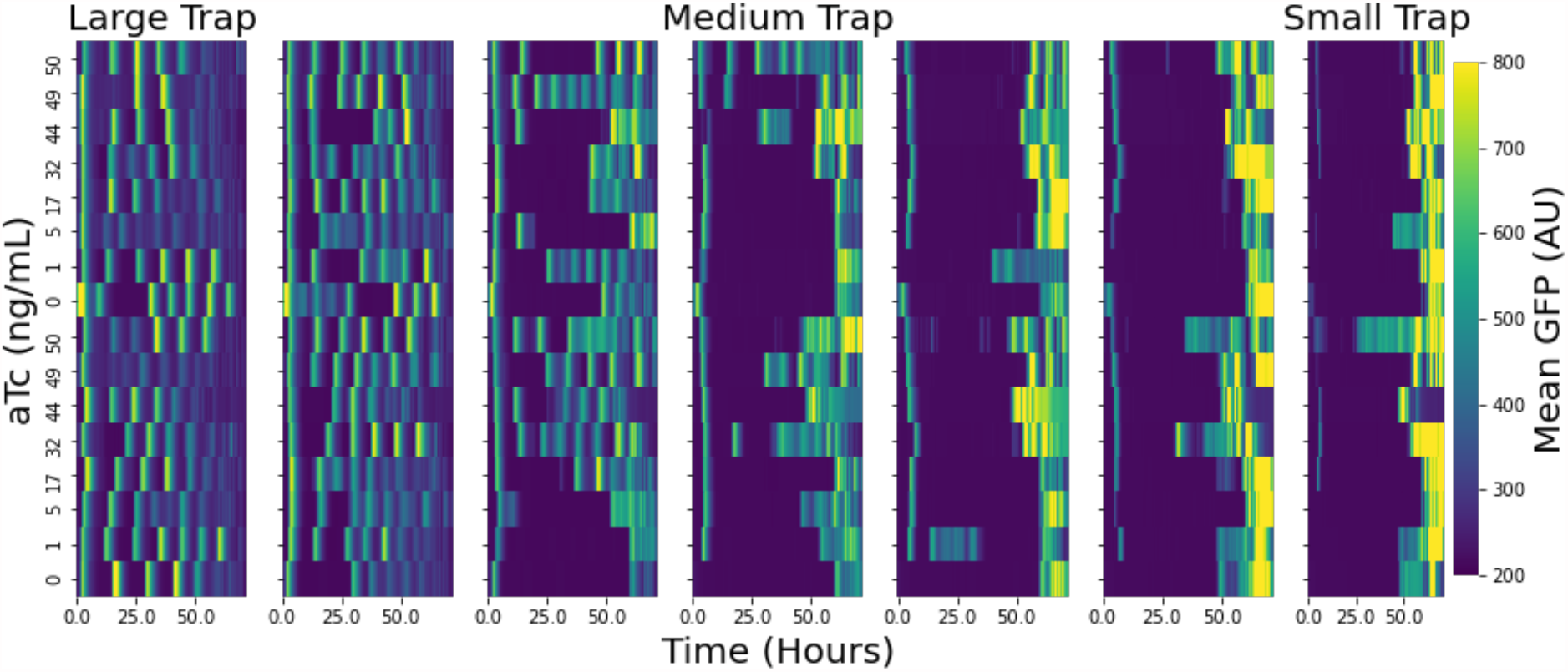
Full single strain microfluidic data set for the TetR-GFP synchronized oscillator library strain D1. Cell trap sizes ranged from 1.0 *\** 10^−2^*mm*^2^ (Large Trap) to 0.16 *\** 10^−2^*mm*^2^ (Small Trap)

**SI Figure 7:**
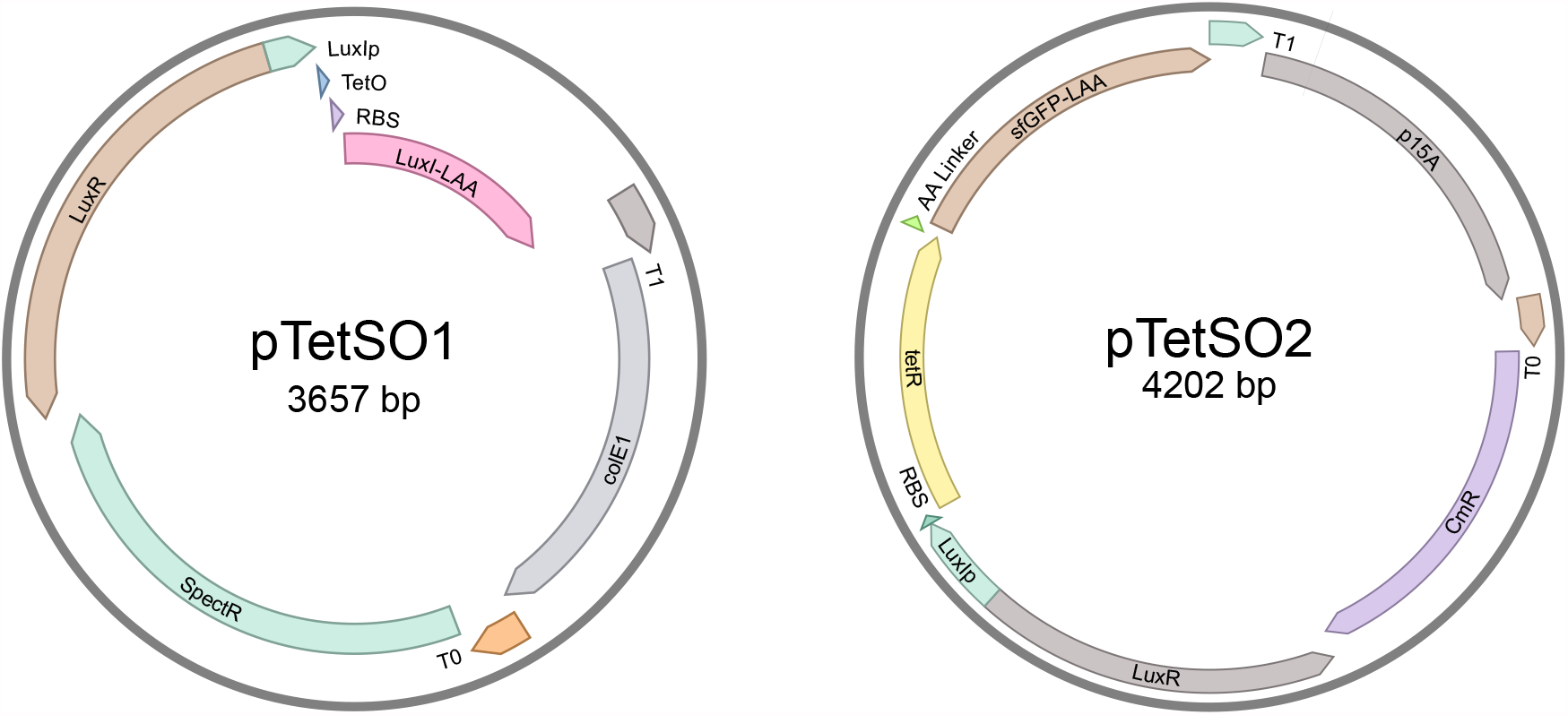
Maps for plasmids used in the TetR-GFP synchronized oscillator.

**SI Figure 8:**
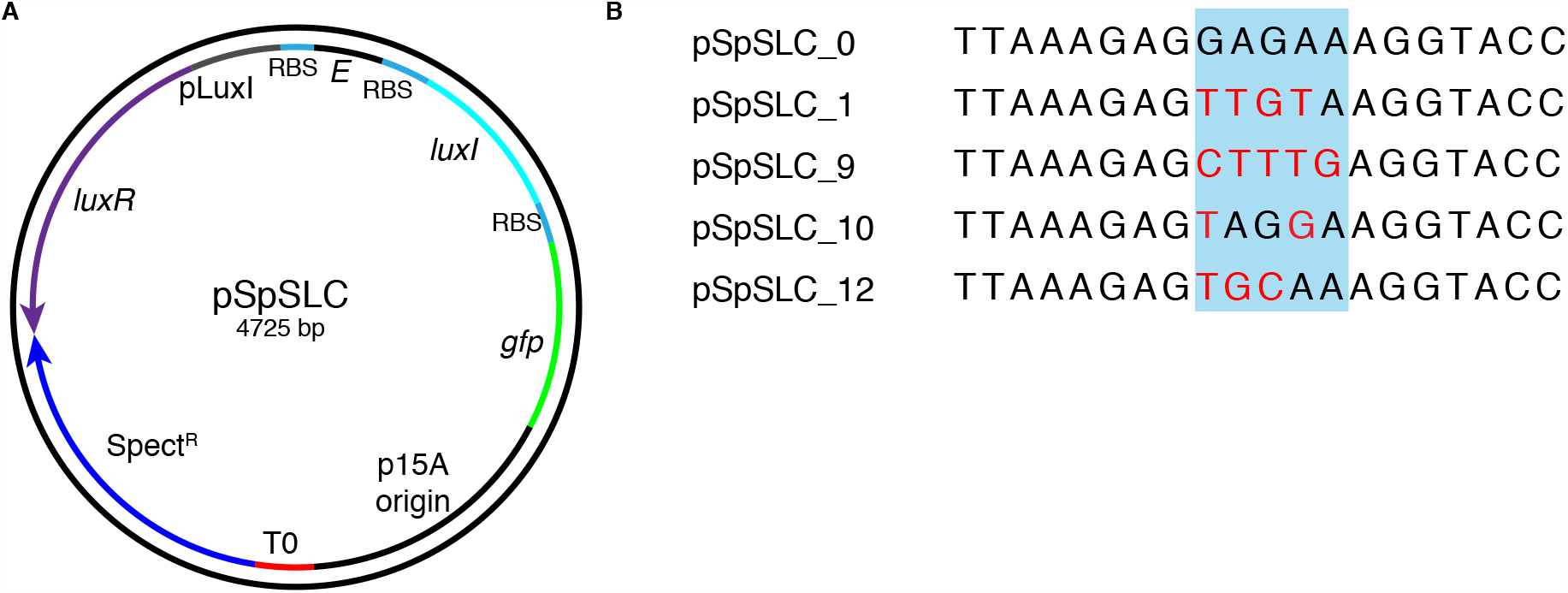
(A) Plasmid map for synchronized lysis circuit. (B) RBS sequences preceding the lysis gene E for selected lysis circuit library members shown in Figure 2. Red base pairs indicate mismatches from original sequence.

**SI Figure 9:**
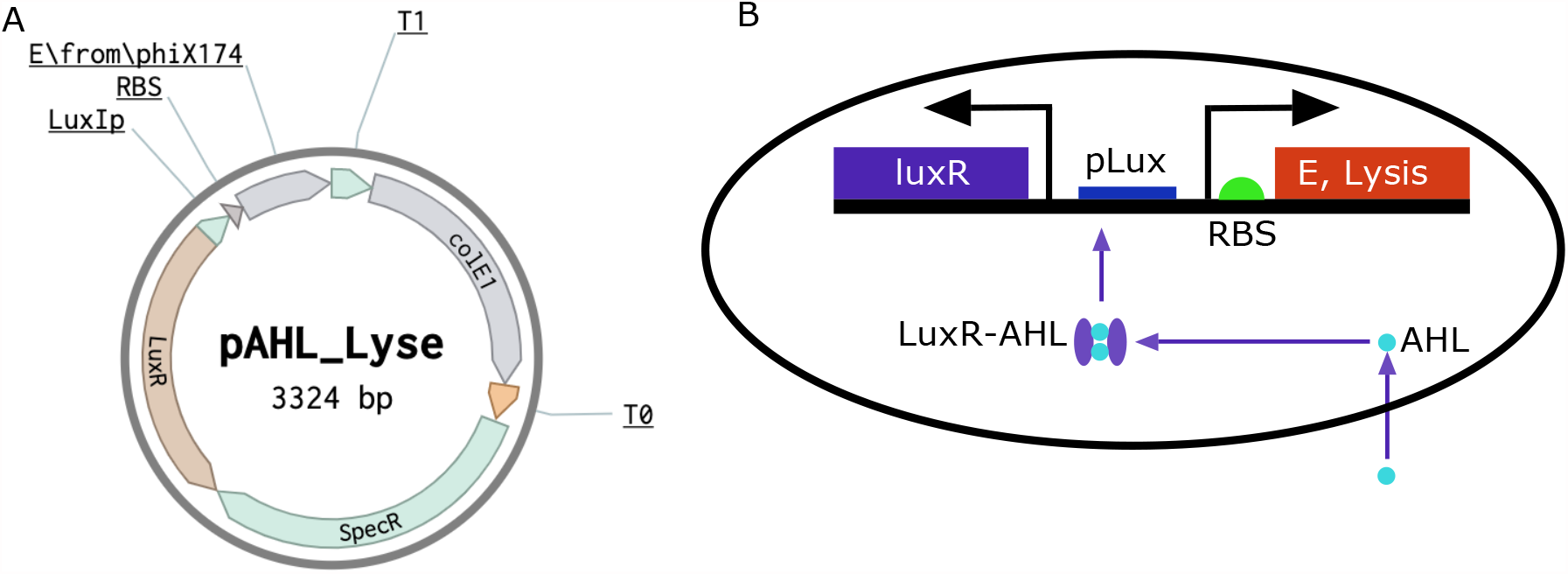
(A) Plasmid map for inducible lysis circuit used to assess RBS strength in (Fig. 3 and SI Fig. 4). (B) Circuit diagram for inducible lysis circuit.

